# Soma-seq links intracellular protein states to transcriptional programs in the human brain

**DOI:** 10.64898/2026.05.28.728561

**Authors:** Carmen Sandoval-Filarsky, Caspar Glock, José Miguel Andrade López, Max Adrian, Vishwa Talati, Oded Foreman, Jeffrey W. Hofmann, Casper C. Hoogenraad, Orit Rozenblatt-Rosen, Brad A. Friedman, Michelle B. Chen

## Abstract

Neurodegenerative disease is largely driven by pathological protein states, which cannot be fully inferred from transcript levels. Yet, joint measurement of cytoplasmic proteins and RNA in single cells from archival human brain tissue remains challenging. Here we present Soma-seq, a method integrating transcriptome and antibody-based intracellular protein measurements in single cells from frozen human brain tissue. Applying Soma-seq to Alzheimer’s disease cortex, we quantified multiple intracellular proteins, including hyperphosphorylated Tau, alongside RNA. Soma-seq resolves a continuous trajectory of pathological progression based on multiplexed protein measurements and reveals associated gene programs. CRISPR perturbation in human iPSC-derived neurons validates Soma-seq-derived candidates and identifies protective factors to Tau aggregation.

## Main

Single cell multiomics of human brain tissue has transformed our ability to dissect the molecular mechanisms of neurodegeneration at cell-type resolution and predict causal regulators of disease^1–4^. However, a central gap remains: most approaches infer cellular states and function from RNA, despite the fact that neurodegenerative pathology is largely defined at the level of proteoforms^5^. Furthermore, RNA and protein abundance are frequently decoupled during brain development, aging, and disease^6–9^, limiting the ability to infer proteomic outcomes from transcriptomic measurements alone. This disconnect is particularly pronounced in neurodegeneration, where disease-defining features arise from post-translationally modified proteoforms, including phosphorylated Tau in Alzheimer’s disease (AD)^10^, α-synuclein in Parkinson’s disease (PD)^11^, and TDP-43 in amyotrophic lateral sclerosis (ALS)^12^. A framework that directly links gene expression to intracellular proteins at single-cell resolution in human brains would enable systematic identification of functionally relevant disease states and their regulators.

Cellular Indexing of Transcriptomes and Epitopes by sequencing (CITE-seq)^13^ enables joint measurement of RNA and protein, but existing implementations have largely been restricted to extracellular epitopes, limiting their ability to interrogate intracellular pathology. Intracellular proteins remain difficult to profile due to fixation-induced disruption of poly(dT)-based reverse transcription and elevated background from nonspecific interactions between oligonucleotide-conjugated antibodies and intracellular components. Recent methods, including ASAP-seq^14^, inCITE-seq^15^, and NEAT-seq^16^ extend multimodal profiling to intranuclear proteins but do not capture cytoplasmic epitopes. Conversely, Phospho-seq^17^ profiles intracellular proteins alongside chromatin accessibility, but lacks matched RNA measurements. Critically, these methods have been largely confined to cell lines, organoids, or fresh peripheral blood, and have not been demonstrated in frozen archival human tissues.

Here, we present Soma-seq, a broadly applicable method for simultaneous profiling of cytoplasmic proteins and RNA in single cells from archival human brain tissues. Soma-seq preserves both transcript and protein integrity, enabling quantitative measurement of intracellular protein abundance with potential for high antibody plexity. Applying Soma-seq to fresh-frozen human Alzheimer’s disease brain tissue, we uncovered novel neuron subtype-specific gene programs associated with protein-defined pathological progression, a subset of which were validated in an iPSC-neuron model of Tau aggregation.

To enable intracellular CITE-seq in archival brain tissue, we leveraged recent advances in isolating intact cell bodies (somas) from frozen tissue via mechanical douncing (1.0-1.5 mm clearance), preserving cytoplasmic content (**Figure 1A-B**)^18^. Whereas prior strategies relied on limited markers to pre-sort cells into binary groups^18^, we developed a marker-agnostic enrichment strategy using phalloidin (a simple counterstain for F-actin) and DAPI to isolate cytoplasm-retaining cells by flow cytometry while preserving cellular diversity (**Extended Figure 1A**). Cells were lightly fixed (1.6% formaldehyde, 10 min) immediately following dissociation to preserve intracellular epitopes. Sequencing of DAPI^+^/phalloidin^+^ populations recovered diverse neuronal and glial populations, including multiple excitatory and inhibitory neuronal subtypes as well as astrocytes, microglia, and oligodendrocytes. While phalloidin preferentially enriches excitatory populations, all major cell classes were retained (**Figure 1C-D**). Transcriptomic complexity was comparable to established human brain snRNA-seq datasets^19–24^, indicating that fixation, staining, and sorting do not substantially compromise RNA quality (**Extended Figure 1B-C**).

**Figure 1.**
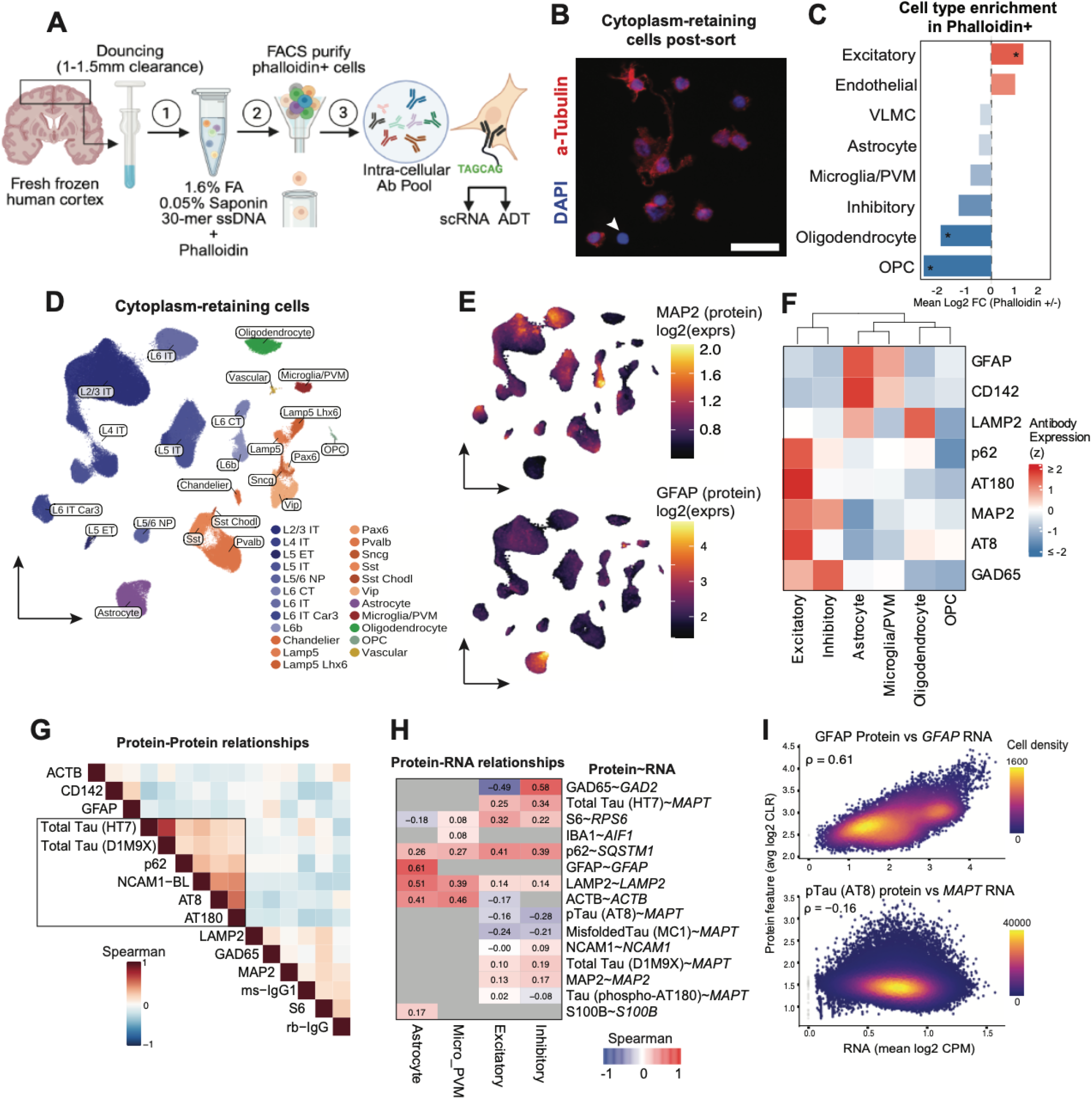
Soma-seq enables simultaneous protein and RNA profiling of neuronal somas from frozen human brain. a) Schematic of workflow: (1) fresh frozen human brain tissue is microdissected and dounced at large clearances followed by light fixation, permeabilization, and blocking. (2) Isolated somas are stained with phalloidin and DAPI, flow enriched, and (3) stained with a Eco-SSB blocked intracellular antibody pool. b) Representative images of phalloidin-positive somas stained with alpha-tubulin. White arrow indicates naked nuclei. Scale bar = 50 um. c) Cell type enrichment in phalloidin+ populations compared to phalloidin- (naked nuclei). *p<0.05, n=3 samples. d) UMAP colored by cell type from FACS-enriched soma-retaining cells. e) Antibody levels of representative intracellular proteins MAP2 (neurons) and GFAP (astrocytes). f) Scaled antibody levels of select cell-type specific epitopes across different neuronal and non-neuronal cell types g) Spearman’s correlation between intracellular proteins. Box highlights correlations between Tau-related proteins. h) Spearman’s correlation between each antibody and its cognate transcript, within each major cell class. Grey indicates epitope is not relevant to a specific cell type. i) Scatterplot of normalized antibody level vs RNA expression at single cell resolution: GFAP ADT vs GFAP RNA (top) in astrocytes, and AT8 ADT vs MAPT RNA (bottom) in excitatory neurons.

A central challenge in intracellular CITE-seq is achieving high signal-to-noise for oligonucleotide-conjugated antibodies in the presence of abundant intracellular nucleic acids and charged proteins. To address this, we combined several techniques into Soma-seq^25,26^: (1) oligo-antibody conjugates were blocked with single-stranded DNA-binding protein (EcoSSB), (2) sorted somas were blocked with random 30-mer ssDNA and (3) lightly permeabilized with 0.05% saponin to increase antibody access without compromising RNA capture (**Figure 1A**). Using a panel of 15 custom-conjugated intracellular antibodies, we demonstrate strong cell-type-specific enrichment of canonical markers, including MAP2 in neurons, GAD65 in inhibitory neurons, and GFAP in astrocytes (**Figure 1E-F**). Control epitopes (ACTB and isotype controls) showed no structured enrichment, confirming low background (**Extended Figure 1D**). We first explored the unique ability of multiplexed proteomic measurements to uncover protein-protein relationships. Antibody-derived tag (ADT) analysis revealed strong concordance between phosphorylated Tau epitopes (AT8, AT180) and weaker association with total Tau (HT7, D1M9X), suggesting selective enrichment of pathological proteoforms. Interestingly, phosphorylated Tau strongly correlated with p62 and NCAM1, linking aggregation to proteostasis and synaptic remodeling pathways (**Figure 1G**). Notably, RNA-protein concordance varied substantially: while markers such as GFAP, GAD65, p62, and LAMP2 showed agreement between modalities, pathological proteoforms exhibited weaker correlations with transcript levels (**Figure 1H-I**), highlighting the limitation of RNA-based inference and motivating direct protein measurement.

Applying Soma-seq to the frontal association cortex from AD (n=11) and control (n=7) donors, we generated a dataset of >200,000 single cells with paired RNA and 13 intracellular protein measurements (**Figure 2A**). We first focused on dissecting hyperphosphorylated Tau - gene relationships using AT8 (pTau Ser202+Thr205), an established marker of Tau pathology in Alzheimer’s disease^27,28^. pTau measured by Soma-seq showed strong concordance with AT8 immunohistochemistry in matched sections, demonstrating quantitative fidelity and good dynamic range (**Figure 2B**). Analysis of neuronal subtypes revealed selective enrichment of pTau in specific excitatory populations, including L5/6 near-projecting (NP), L2/3 and L5 intratelencephalic (IT) neurons, suggesting subtype-specific vulnerability (**Figure 2C**). Differential expression analysis (DEA) between pTau high vs. low populations identified dysregulation of pathways related to microtubule dynamics, stress response, synaptic function, and proteostasis, including genes involved in protein folding, lysosomal trafficking, and ubiquitin-mediated degradation (**Figure 2D-F, Extended Figure 2A-B**), largely consistent with priorpTau-enriched flow studies^18^ (**Extended Figure 3A-D**). While many transcriptional changes were convergent across neuronal subtypes, a subset of high effect-size DEGs exhibited layer-specific regulation, indicating both shared and context-dependent responses to pathology (**Figure 2E, Extended Figure 2C**). These results demonstrate that subsetting cell states by protein pathology reveals distinct transcriptional programs not captured by transcriptome alone.

**Figure 2.**
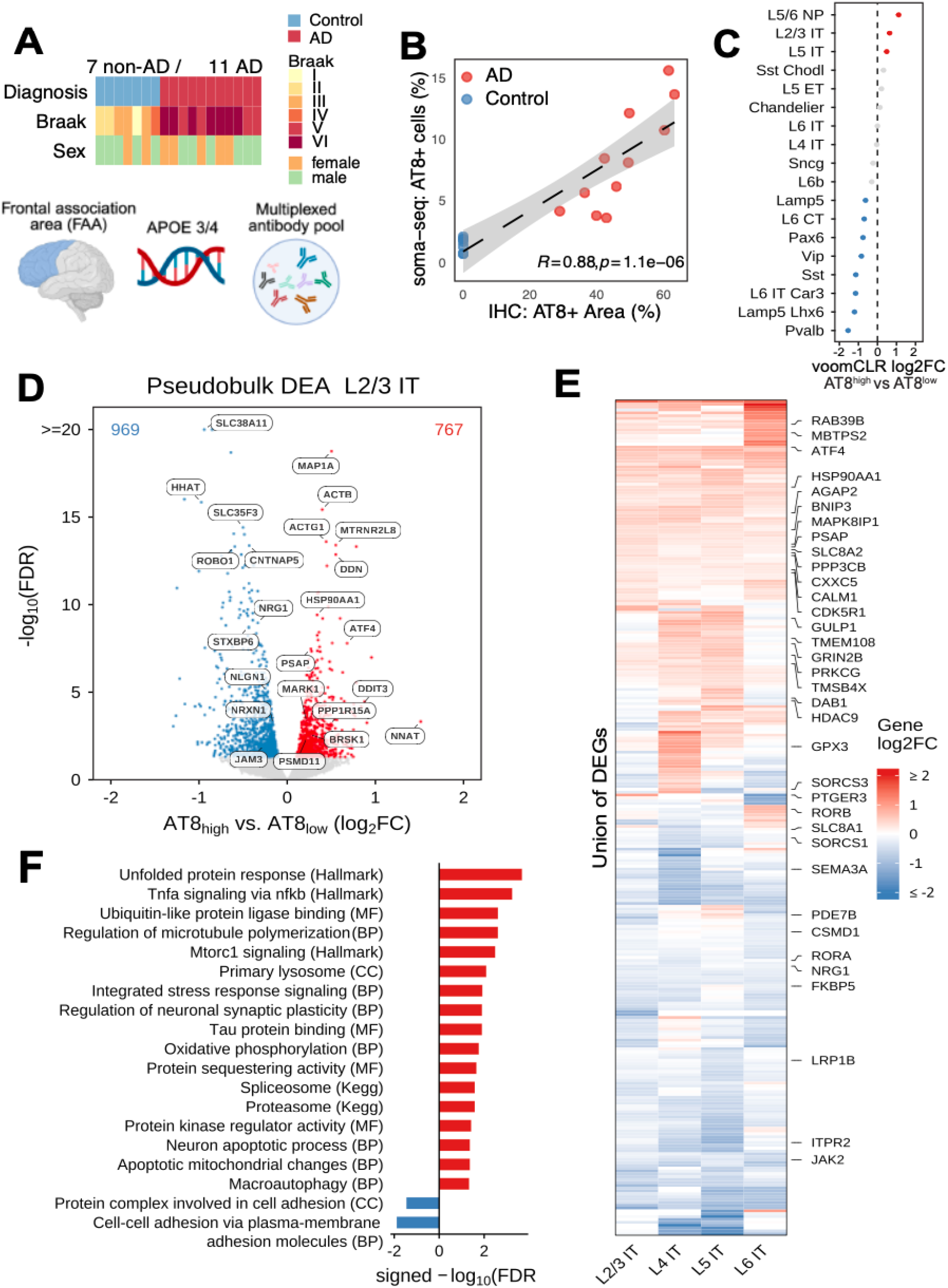
Soma-seq of human AD tissue reveals hyperphosphorylated Tau-specific pathways. a) Study design: Frontal association area (FAA) from 7 non-AD (Braak I-IV) and 11 AD (Braak V-VI) donors were profiled via Soma-seq. b) R-squared comparing single cell AT8 detection by Soma-seq (% of all cells) vs gold-standard IHC (% of AT8 area) in matched sections (n=7 non-AD, 11 AD). c) voomCLR log2 fold-change in neuron subtype composition for AT8-high vs AT8-low neurons. Positive values indicate relative enrichment among AT8-high neurons, while negative values indicate relative enrichment among AT8-low neurons. Red and blue dots indicate neuron subtypes with significant enrichment in AT8-high and AT8-low neurons, respectively. FDR <= 0.05. d) Volcano plot of donor-level pseudobulked differentially expressed genes specifically in L2/3 IT neurons. Red and blue dots indicate genes significantly up- and downregulated, respectively, in AT8 high neurons. FDR ≤ 0.05. Genes of interest are labeled. e) Log2 fold change of the union of top differentially expressed genes across neuron layers. Genes of interest are labeled. f) Pathway enrichment of differentially expressed genes in L2/3 IT neurons (AT8 high vs AT8 low).

To capture the continuous nature of neurodegenerative progression^29,30^, we leveraged both the plexity and quantitative nature of protein measurements provided by Soma-seq rather than imposing binary thresholds on single ADTs. We first constructed metacells^31^ with L2/3 IT neurons (the most abundantly captured neuron-subtype) in the ADT-space using all pathologically-relevant proteins, reducing noise and increasing biological interpretability. This revealed two divergent trajectories: (1) a non-pathological state characterized by high total Tau and low pTau (AT8) present in both AD and control samples, and (2) a pathology-associated trajectory defined by both elevated Tau and pTau, restricted to AD samples (**Figure 3A-B**). Ordering metacells along this axis reveals a continuous protein-based disease trajectory, as supported by the graded expression of other pathologically-associated proteins such as pTau Thr231^32^ (AT180) and p62/SQSTM1^33^. Using this ordering, we applied consensus non-negative matrix factorization (cNMF^34^) to identify gene programs that co-vary along this pathology trajectory (Methods). Modules associated with lysosomal function (M1) and metabolic processes (M7) increased with pathology, whereas synaptic and neuronal projection programs were progressively downregulated (M2, 4, 5), consistent with recent findings^18,22,23^ (**Figure 3C**). These results demonstrate that proteoform-defined trajectories enabled by Soma-seq can provide a principled framework for linking transcriptional programs to progressive cellular disease states.

**Figure 3.**
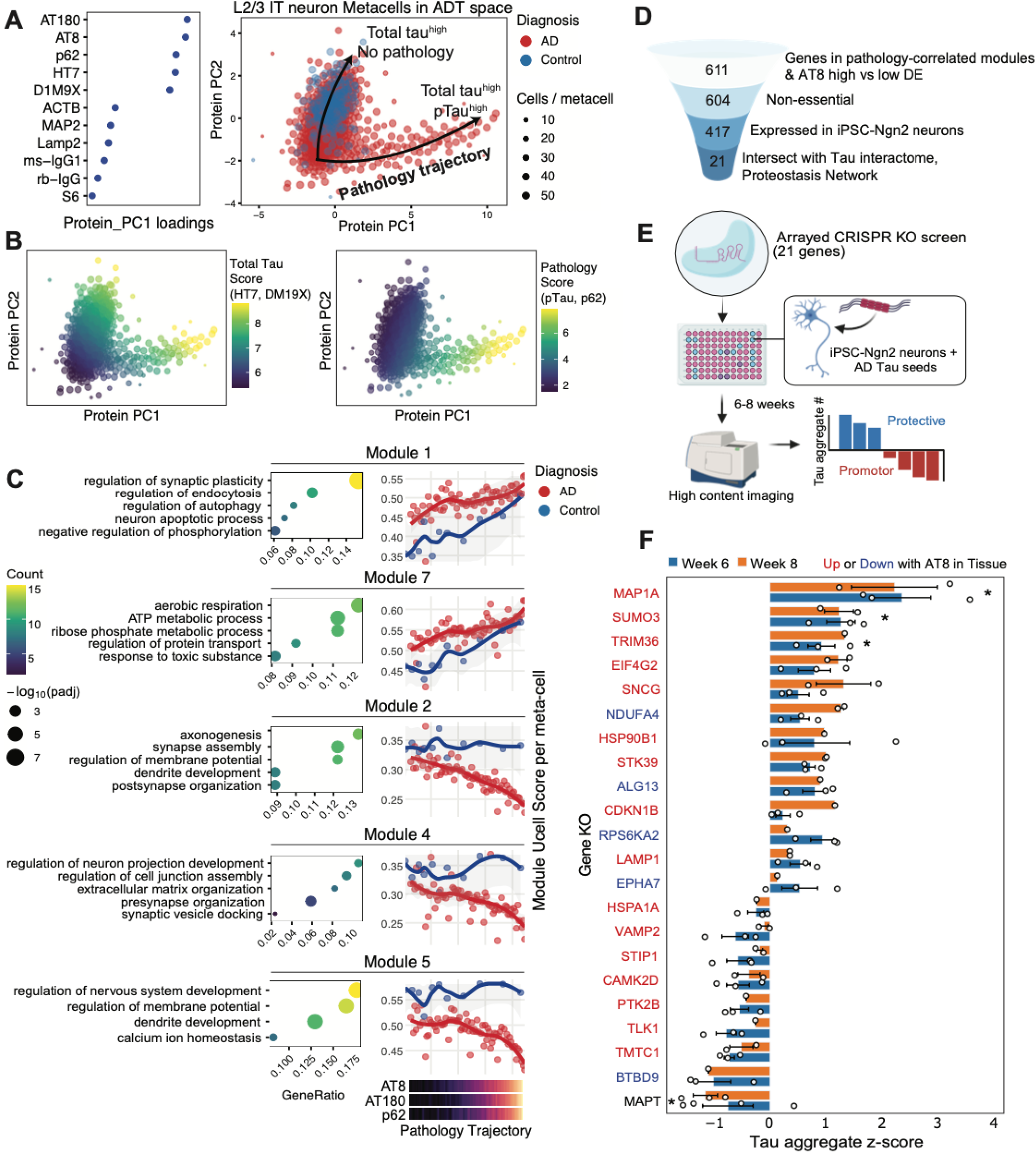
Multiplexed protein measurements resolve a continuous pathology trajectory and associated gene programs. a) PC1 loadings of all neuron-relevant antibodies in ADT feature space. PCA plot of L/23 excitatory neuron metacells constructed via KNN aggregation in ADT feature space. Two potential trajectories (pathological vs non-pathological) are labeled by arrows. b) PCA of metacells colored by Total Tau score (normalized expression of HT7, D1M9X combined) and Pathology Score (normalized expression of AT8, AT180, p62 combined) on the ADT-only representation c) (Right) UCell scores of gene modules increasing (M1, M7) or decreasing (M2, M4, M5) along the pathology trajectory defined in b) (metacells ranked by averaged PC1 coordinate). Bottom legend indicates scaled expression of pathology-associated proteins AT8, AT180 and p62. (Left) Select top GO ontology terms enriched in the top loading genes in gene modules d) Screening funnel: pathology-associated genes discovered by Soma-seq are further filtered by essentiality, expression in iPSC neurons, and biological relevance. e) Screening schematic: CRISPR RNPs of 21 gene candidates are delivered via electroporation into iPSC-NGN2-neurons seeded with human AD brain-derived Tau seeds. High content imaging is performed after 6 and 8 weeks. f) Z-scores of aggregate counts across each CRISPR perturbation relative to controls at 6 and 8 weeks post-seeding. Each point indicates an independent well. Bars depict mean and error bars depict standard deviation. * indicates FDR<0.05 and |z-score|>1 after a one-sample two-tailed Student’s t-test.

Finally, we validated candidate regulators of Tau pathology identified by Soma-seq using arrayed CRISPR knockout screening in human iPSC-derived neurons^35^ with endogenously-tagged Dendra-Tau^36^ (Andrade Lopez et al., submitted) (**Figure 3D**). Day 7 neurons were edited with Cas9 ribonucleoproteins targeting 21 candidate genes and exposed to AD brain-derived Tau seeds^37^ for 6 to 8 weeks, followed by high-content imaging of aggregate formation (**Figure 3E**). As expected, MAPT knockout reduced the development of Tau aggregates. Additionally, we identified three genes - *TRIM36* (a microtubule-associated E3 ligase)^38^, *MAP1A* (a regulator of microtubule dynamics)^39,40^, and *SUMO3* (a small ubiquitin-like modifier) ^41^ - that suppress seed-mediated Tau aggregation (**Figure 3F, Extended Figure 4**). These genes were progressively upregulated along AT8 pathology trajectories in AD brains, suggesting the emergence of a compensatory protective program linking microtubule stabilization with SUMO/ubiquitin-associated proteostasis. Such coordinated regulation may limit the pool of seed-competent Tau by promoting microtubule association and enhancing aggregate handling or clearance.

Finally, we demonstrate the ability of Soma-seq data to impute protein expression levels on existing scRNA-seq datasets lacking direct protein measurements. Using our atlas as a reference, we successfully imputed AT8 levels in L2/3 IT neurons from the Otero et al dataset^18^, which measured only RNA but with known protein state labels (AT8 high vs AT8 low) acting as a ground truth (**Extended Figure 3E-F**). This establishes a framework for the projection of intracellular protein states to unimodal scRNA-seq datasets, allowing the identification of transcriptional states concordant with cellular proteinopathy.

Future work includes further increasing the plexity of antibodies to better infer function, particularly pathologically meaningful epitopes such as proteostasis regulators, mitochondrial/oxidative stress and cell death markers. Lastly, Soma-seq can be applied to other neurodegenerative diseases where pathology is defined by specific protein-level states (e.g. post-translational modifications, conformations, cleavage products), such as alpha-synuclein (pS129) in PD, TDP-43 (pS409/410) in ALS, and polyglutamine-expanded HTT in Huntington’s disease. Altogether, Soma-seq enables discovery and functional validation of genetic regulators of protein pathology, and offers a platform for resolving proteoform-defined cell states.

## Experimental Methods

### Sample Selection

Frozen postmortem human brain tissue was obtained from 18 donors (11 Alzheimer’s disease cases, 7 age-matched controls). AD cases were Braak stage V-VI, while controls were Braak stage I-IV. All donors were APOE ε3/ε4 heterozygotes. Tissue samples were dissected from the frontal association cortex, spanning all cortical layers but excluding white matter.

### Antibody-oligonucleotide Conjugation

Custom antibody-oligonucleotide conjugates were generated using IEDDA click chemistry. Briefly, amino-C12 modified oligonucleotides (100 nmol scale, IDT) were first purified by ethanol precipitation, then conjugated to trans-cyclooctene (TCO) by reacting with TCO-PEG4-NHS (Click Chemistry Tools, 100 mM in DMSO) in borate-buffered saline (BBS) pH 8.5 for 30 minutes at room temperature. TCO-modified oligonucleotides were desalted using Bio-Rad Micro Bio-Spin P-6 columns and adjusted to 100 μM in 1× BBS. BSA-free antibodies (50 μg each) were buffer-exchanged into 1x BBS using Amicon Ultra-0.5 30 kDa MWCO filters and conjugated to methyltetrazine (mTz) by reacting with mTz-PEG4-NHS (Click Chemistry Tools, 2 mM final concentration) for 30 minutes at room temperature. Excess NHS groups were quenched with 1M glycine pH 8.5, and mTz-labeled antibodies were purified using 30 kDa MWCO filters. For final conjugation, mTz-antibodies were reacted with TCO-oligonucleotides at a ratio of 30 pmol oligo per 1 μg antibody overnight at 4C. Residual tetrazine sites were quenched with TCO-PEG4-glycine (10 mM). Antibody-oligonucleotide conjugates were purified by repeated ammonium sulfate precipitation (40% saturated ammonium sulfate, overnight at 4°C followed by 2 additional 1-hour cycles) to remove unconjugated oligonucleotides, followed by 3 washes with PBS using Amicon Ultra-0.5 50 kDa MWCO filters. Conjugate purity was verified by 4-15% TGX Stain-Free protein gels and 4% E-gel EX agarose gels. Final conjugates were adjusted to 0.5 μg/μL in PBS containing 1 μg/μL BSA and 0.06% sodium azide and stored at 4°C. The complete antibody panel and barcode information is provided in Supplementary Table X.

### Tissue Dissociation and Fixation

30 mg of tissue per donor were processed together. Homogenization was performed as described in Otero et al, 2022. Briefly, tissue was cut into small pieces on a glass petri dish chilled on ice. Gentle mechanical dissociation was performed using an 8 ml glass Potter-Elvehjem homogenizer with PFTE pestle (1-1.5 mm clearance) in chilled soma-prep buffer (2 mL per 100 mg tissue). Soma-prep buffer contained 10 mM Tris-HCl pH 8.0, 5 mM MgCl_2_, 25 mM KCl, 250 mM sucrose, 0.5× protease inhibitor cocktail (cOmplete, Roche), 1 μM DTT, 2 U/mL DNase I, and 1 U/μL RNase inhibitor (Protector, Roche), 1x PBS, and was prepared at 2 mL per 100 mg tissue. 4-5 gentle strokes were used until large tissue chunks were not observed. Cell suspension was strained through a 100uM mesh filter, then a 50uM mesh filter, and split evenly between 10 5% BSA-coated 1.5 ml Protein-lo bind tubes. Dissociated tissue was immediately fixed by adding 32% formaldehyde to a final concentration of 1.6% and incubating for 10 minutes at room temperature. Fixation was quenched with 2.5 M glycine to a final concentration of 0.125 M. Fixed cells were pelleted by centrifuging at 250xg for 5 minutes in 5% BSA-coated 1.5 mL Protein lo-bind tubes. Pellets were resuspended in 1ml staining buffer (1% BSA, 0.05% saponin, 5 mM MgCl_2_, 0.05 U/ul Protector RNase Inhibitor). Cells were stained with phalloidin (150 nM final concentration) and DAPI (1 ug/ml). Cells were incubated with phalloidin/DAPI for 30 minutes at room temperature on a rotator. Following staining, cells were washed three times in wash buffer (3% BSA, 5 mM MgCl_2_ in PBS) by centrifugating at 450xg for 5 minutes.

### Flow Cytometry and Sorting

Phalloidin/DAPI-stained cells were sorted on a Sony MA900 fluorescence-activated cell sorter by first selecting DAPI-positive events to discard debris and other non-nucleated events. Nucleated events were then gated on a phalloidin vs SSC. Approximately 1.0×10^6^ cells were recovered after sorting, with a target recovery of 50,000 cells per donor.

### Antibody blocking and cell staining

The antibody pool was incubated with single-stranded DNA binding protein (SSB) to block oligonucleotide barcodes and reduce unspecific binding, following the NEAT-seq protocol. SSB was added at a ratio of 10 μg SSB per 1 μg of total antibody in 1x NEB buffer. Sorted cells were pelleted by centrifugation, resuspended in 50ul of Staining Buffer, and stained with 50 ul of the pre-blocked custom CITE-seq antibody cocktail. Antibodies were diluted to a final concentration of 5 μg/mL. Cells were incubated for 1 hour at 4C on a rotator and washed three times in Wash Buffer (3% BSA, 5mM MgCl_2_ in PBS) by centrifugating at 450xg for 5 minutes. After the last wash, the pellet was resuspended in 30 ul of PBS without Ca/Mg. Cells were counted on a hemocytometer by diluting a 3ul aliquot of the cell suspension in PBS.

### Single-cell capture and cDNA synthesis

Cells were loaded onto the 10x Genomics Chromium Controller using the Chromium Single Cell 3’ Gene Expression HT v3.1 kit. Cells were loaded at approximately 115,000 cells per channel across 7 channels, targeting 60,000 cells recovered per channel (20,000 recovered cells per donor). Following cell capture and barcoding, reverse transcription, cDNA amplification and SPRI bead cleanup were performed according to the 10x Genomics protocol. During cDNA amplification, CITE-seq Additive Primer (200nM final concentration) was spiked into the PCR master mix to enable co-amplification of ADT-derived sequences alongside gene expression cDNA, following the standard CITE-seq protocol. Following amplification, cDNA product was size-separated using a double-sided SPRI bead selection to isolate small (<300 bp) and large (>300 bp) size fractions, following the 10x Genomics Feature Barcode protocol. The small fraction, enriched for ADT-derived sequences, was used for ADT library preparation. The large fraction was used for gene expression (GEX) library preparation.

### Library preparation and sequencing

Gene expression (GEX) libraries were prepared according to the standard 10x Genomics Chromium Single Cell 3’ v3.1 protocol. ADT libraries were prepared from the ADT cDNA fraction following the standard CITE-seq guidelines. Briefly, ∼50ng of ADT cDNA was indexed with ADT-specific primers using NEBNext with 17 PCR cycles. ADT and GEX libraries were purified using SPRI beads. Library quality and concentration were assessed using Qubit (High Sensitivity dsDNA kit) and Agilent TapeStation HS D1000. GEX libraries showed expected size distributions around 460 bp. ADT libraries showed expected size distributions around 220-231 bp. Libraries were pooled and sequenced on an Illumina NovaSeq platform. GEX libraries were sequenced using standard 10x Genomics read configurations (Read 1: 28 bp, i7 index: 10 bp, i5 index: 10 bp, Read 2: 90 bp). Target sequencing depth was 25,000 reads per cell for GEX libraries and 5,000 reads per cell for ADT libraries.

### CRISPR screening in IPSC-neurons

Human iPSCs (IP11N, Alstem) containing a Doxycycline-inducible *NGN2* transgene were engineered in-house to include an endogenous N-terminal Dendra2-tag on wild-type Tau and were differentiated into iNeurons according to Shan et al., 2024. In brief, iPSCs were maintained on iMatrix-coated plates in mTeSR Plus. At 80% confluence, iPSCs were dissociated with Accutase and plated at 20-40K cells/cm^2^ in Induction Media (DMEM/F12, N2, B27, Glutamax, NEAA) supplemented with SB431542, XAV-939, Noggin, and Doxycycline (3μg/ml) to initiate *NGN2* expression. Following 4 days of induction, cells were transitioned to Maturation Media on Day 5 with 5 μM AraC and 2 μg/ml Doxycycline for 2 days to remove non-neuronal cells. On Day 7, iNeurons were dissociated with Accutase and cryopreserved.

DIV7 iNeurons were edited by nucleofection of Cas9 ribonucleoprotein (RNP) complexes targeting 21candidate genes. RNPs were assembled by combining IDT Alt-R S.p. Cas9 protein with pre-designed synthetic guide RNAs (Gene Knockout Kits, EditCo) at a 3:1 guide-to-Cas9 molar ratio. RNPs were delivered into neurons using the Lonza 4D-Nucleofector system in 96-well Nucleocuvette plates (program CV110) . Neurons were then seeded into 96-well PhenoPlates, cyclic olefin bottom (Revvity) at 100-150K cells/cm^2^ onto PDL and iMatrix-coated plates in Maturation Media (Neurobasal, BDNF, GDNF, Ascorbic Acid, cAMP, TGF-β3). Doxycycline (2 μg/ml) and DAPT were supplemented during the first two weeks of maturation to ensure stable neuronal identity.

Tau aggregation was induced by seeding iNeurons at DIV14 with Alzheimer’s disease patient-derived tau (ADtau) fibrils extracted from the human cerebral cortex. ADtau fibril concentration was determined by total tau ELISA and added at a final concentration ranging from 50-110 pg/mL. Cultures were maintained for 6-8 weeks, after which Tau aggregation was assessed in live neurons by high-content imaging on an ImageXpress HT.ai confocal microscope (Molecular Devices) using an Apo 40X 1.15 NA LWD water immersion objective (Nikon Microsystems) capturing the entire well surface. After TopHat filtering (7px) Tau aggregates were defined as structures approx 0.2-5 µm wide with an intensity of >1500 above local background, an ellipticity of >1.5 and an area >100 nm^2^. Rounder, dimmer structures 1-5 um wide were excluded as Wallerian degradation blebs commonly found in neuronal cultures. Aggregate counts and total Dendra2 area (as a proxy for cell density) were measured using the Custom Module Editor in MetaXpress software (Molecular Devices).

## Computational Methods

### Soma-seq data pre-processing and quality control

Raw sequencing data were demultiplexed to FASTQ files using Cell Ranger *mkfastq* (v7, 10x Genomics). Reads were aligned to the human reference genome (GRCh38) with intronic reads included, and gene expression was quantified as unique molecular identifier (UMI) counts using Cell Ranger *count* (v7, 10x Genomics). Ambient RNA contamination was removed by processing all samples with CellBender^42^ (v0.3) using the *raw_feature_bc_matrix*.*h5* output from Cell Ranger as input, with default parameters and excluding antibody-derived tag (ADT) counts. Barcodes classified as somata by CellBender, together with the corresponding decontaminated count matrices, were used for downstream analyses.

Per-soma nuclear fraction scores were computed using DropletQC^43^ applied to the Cell Ranger output. Cells were required to pass the following initial quality-control thresholds: at least 200 detected genes, at least 400 UMIs, and at most 50% mitochondrial reads. Donor pools were genotype-demultiplexed with Souporcell^44^ using known donor genotypes. Only singlets were retained, and somata lacking a donor assignment were excluded. Additional quality-control thresholds for ADT measurements were defined by applying DropletUtils *cleanTagCounts*^*45*^ to all somata. Cells with elevated IgG signal, defined as greater than 5 median absolute deviations above the median, were removed. Cells with zero estimated ambient contamination were also removed. Within each donor pool, doublets were identified using scDblFinder^46^, run separately for each donor with default parameters.

For cell-type annotation, labels were transferred from the SEA-AD snRNA-seq reference dataset^47^ using Seurat (v5) *FindTransferAnchors* and *TransferData*^*48*^.

### Soma-seq data integration and cell type interpretation

Data integration was performed using Harmony as implemented in Seurat (v5). Count matrices were loaded into a Seurat object, split by donor, and processed using default parameters unless otherwise stated. The following functions were applied in order: *NormalizeData, FindVariableFeatures* with *nfeatures = 3000, JoinLayers, ScaleData, RunPCA*, and *RunHarmony* with *group*.*by*.*vars = “case_id”* to integrate across donors. Cells were clustered using *FindNeighbors* with *reduction = “harmony”* and *dims = 1:50*, followed by FindClusters with resolution = 3.2. For each cluster, the fraction of predicted doublets identified by scDblFinder and the percentage of transferred cell-type labels were computed.

Broad cell types were assigned to clusters based on the transferred cell-type label distribution and expression of canonical marker genes, including AQP4 for astrocytes, CLDN5 for endothelial cells, SNAP25 for pan-neuronal cells, SLC17A7 for excitatory neurons, GAD2 for inhibitory neurons, CX3CR1 for microglia and perivascular macrophages, PLP1 for oligodendrocytes, and PDGFRA for oligodendrocyte precursor cells.

Clusters with elevated predicted doublet fractions and co-expression of marker genes from multiple broad cell types were annotated as doublets. Clusters were annotated as low quality when they exhibited poor QC profiles, including high mitochondrial read fractions, low numbers of detected genes or UMIs, and high nuclear fraction scores. Doublet and low-quality clusters were excluded from downstream analyses.

Dataset refinement was performed iteratively. After removing low-quality somata and predicted doublets, the integration, clustering, and annotation workflow described above was repeated. The dataset was then partitioned into neuronal and non-neuronal compartments, and the neuronal compartment was further subdivided into excitatory and inhibitory neurons. At each partitioning step, cells were re-integrated and subclustered at high resolution, followed by re-annotation using transferred labels and marker gene expression. Different numbers of principal components were used for integration depending on the partition, with 40 principal components used for analyses including all neurons and 30 principal components used for non-neuronal, excitatory neuronal, and inhibitory neuronal partitions. After the final iteration, all remaining low-quality somata and predicted doublets were removed, and the retained somata were re-integrated and annotated for downstream analyses. In a final step aimed at higher-resolution neuronal subtype classification, excitatory and inhibitory neurons were isolated and analyzed separately to resolve subtypes within each neuronal class. For each high-resolution subcluster, the final neuronal subtype was assigned by majority vote, defined as the most frequently transferred label among somata in that subcluster. When neuronal classes were analyzed in isolation, the SEA-AD snRNA-seq reference was subset to the corresponding neuronal class before label transfer.

### ADT count normalization

ADT counts were normalized following the OSCA “Integrating with protein abundance” workflow (https://bioconductor.org/books/3.22/OSCA/). Baseline ADT abundances were estimated from the ADT count matrix using *ambientProfileBimodal*. Size factors were then computed with *medianSizeFactors* using the baseline profile as the reference.

### AT8_high_ vs AT8_low_ differential expression analysis

For each excitatory neuron subtype, AT8_high_ neurons were defined as having normalized AT8 levels greater than 2.5 median absolute deviations above the subtype-specific median. All remaining neurons were classified as AT8_low_. Pseudo-bulk expression profiles were generated by aggregating UMI counts across somata sharing the same neuronal subtype label, AT8 levels status, and donor identity. Differential expression was performed separately within each neuronal subtype, excluding pseudo-bulk samples derived from fewer than 50 somata. Raw counts were analyzed using edgeR. Lowly expressed genes were removed using filterByExpr and library sizes were normalized with *calcNormFactors* using the *TMMwsp* method, which is designed to better accommodate sparse count profiles with many zeros. Models were fit using the design formula *∼ case_id + adt_status*, where *case_id* encodes donor identity, and *estimateDisp, glmQLFit*, and *glmQLFTest* were used to test for differences between AT8_high_ and AT8_low_ pseudo-bulk samples. Genes with a false discovery rate at most 0.05 were considered differentially expressed.

### Gene set enrichment analysis

Gene set enrichment analysis was performed on pseudo-bulk differential expression results using clusterProfiler gseGO and GSEA. Genes were ranked by signed -log_10_(P value), with the sign taken from the direction of effect in the differential expression analysis. Gene sets with sizes between 10 and 500 were tested, including Gene Ontology terms and Hallmark and KEGG gene sets obtained via msigdbr (https://cran.r-project.org/package=msigdbr). For Fig. 2, significant terms associated with transcriptional changes in AT8_high_ L2/3 IT neurons were visualized. For Extended Data Fig. 2B, visualization followed a consistent, unbiased selection procedure across neuronal subtypes. Within each subtype, terms were ranked by signed -log_10_(adjusted P value) and categorized as shared if they were significantly enriched or de-enriched in more than one subtype in the same direction, or exclusive if they were significant in only a single subtype. For each subtype, up to three top enriched and up to three top de-enriched terms were selected per ontology from both the shared and exclusive sets, when available.

### Re-analysis of Otero-Garcia et al. 2022

FASTQ files were downloaded from SRP190819. Gene expression counts were generated using Cell Ranger count, and ambient RNA contamination was removed using CellBender as described above. Low-quality cells were removed using sample-specific thresholds. Cells were excluded if they met any of the following criteria: UMI counts below 200 or more than 5 median absolute deviations (MAD) below the sample median, detected genes below 200 or more than 5 MAD below the sample median, percentage of counts in the top 20 genes more than 5 MAD above the sample median, or mitochondrial read fraction above 20% or more than 3 MAD above the sample median. Doublets were predicted using scDblFinder as described above, and cell-type labels were transferred from the SEA-AD reference as described above. Cell-type annotation was based on the same marker genes used in the primary analysis.

Dataset cleaning followed the same iterative strategy described above, with small variations. Samples were integrated using Harmony. The Seurat object was split by sample and processed using NormalizeData, FindVariableFeatures with 3,000 features, ScaleData, RunPCA, IntegrateLayers with method = HarmonyIntegration and npcs = 30, RunUMAP with reduction set to “harmony”, FindNeighbors with reduction “harmony” and dimensions 1–30, FindClusters with resolution 3, and JoinLayers. After the first integration round, only excitatory and inhibitory neurons were retained. Two additional rounds of integration were performed, after which the dataset was split by neuronal class and subclusters were assigned to excitatory and inhibitory neurons, respectively, as described above.

Pseudo-bulk expression profiles were generated by aggregating UMI counts across cells sharing the same neuronal subtype label, AT8 status, and donor identity. Differential expression was performed separately within each neuronal subtype, excluding pseudo-bulk samples derived from fewer than 10 cells. Raw counts were analyzed using edgeR. Lowly expressed genes were removed using filterByExpr, and library sizes were normalized with calcNormFactors using the TMMwsp method. Models were fit using the design formula ∼ subject + facs, where subject encodes donor identity and facs encodes AT8 status, and estimateDisp, glmQLFit, and glmQLFTest were used to test for differences between AT8-positive and AT8-negative pseudo-bulk samples. Genes with a false discovery rate at most 0.05 were considered differentially expressed.

### Differential composition analysis

Differential composition between AT8-positive and AT8-negative neurons in Otero-Garcia et al. and between AT8_high_ and AT8_low_ neurons in Soma-seq was tested using voomCLR following the developers’ vignette, using a random-intercept model to account for donor-level correlation^49^. voomCLR applies a centered log-ratio transformation to count data and estimates observation-level heteroscedasticity weights for downstream statistical testing.

For Soma-seq, AT8_high_ neurons were defined as having normalized AT8 levels greater than 2.5 median absolute deviations above the neuron median, and all remaining neurons were classified as AT8_low_. Cell types with an adjusted P value at most 0.05 were considered to exhibit significant compositional differences.

### ADT-based metacell construction

To define protein-resolved cellular states, we constructed metacells directly in the antibody-derived tag (ADT) feature space. Median normalized ADT measurements were used to perform principal component analysis (PCA) using *prcomp* restricted to select neuronal-associated markers (e.g., AT8, AT180, p62, D1M9X, HT7, NCAM1-BL, MAP2). Metacells were generated using a greedy aggregation strategy in ADT PCA space. For each unassigned cell, a seed cell was selected and grouped with its nearest neighbors within the k-nearest neighbor graph, restricting to unassigned cells and aggregating up to a fixed number of cells per metacell (typically 50). This process was iterated until all cells were assigned, yielding non-overlapping metacells that preserve local structure in protein space while reducing single-cell noise. Metacell construction was performed separately for disease and control samples to avoid cross-condition mixing. For downstream analysis, metacell-level features were computed by averaging ADT expression and derived scores (including PC1-based pathology scores) across constituent cells. This approach enables denoising and robust identification of protein-defined cellular states while preserving continuous variation in intracellular proteoforms.

### cNMF and correlations with pathology trajectory

Cells were first subsetted to only those containing low to high pathology (non-zero). A nearest-neighbor graph was constructed from the ADT embedding using all retained dimensions (k = 30), and Leiden clustering was iterated over a range of resolutions (0.1 to 1.0). Clusters were visualized across resolutions in the ADT PCA embedding, and the pathology-relevant population was defined as the cluster that scored consistently high on pathological protein (AT8, AT180, p62) expression. Metacells consisting of these subsetted “pathology-positive” cells were ranked along a continuous pathology axis defined by the mean PC1 of constituent cells, as PC1 is defined primarily by pathologically relevant neuronal protein features. Consensus negative matrix factorization (cNMF) was performed on L2/3 IT neurons using the top 5,000 highly variable genes, and the average UCell score per module per metacell was calculated and visualized by pathology trajectory ranking (ranking of position along PC1). For each module, genes were ranked by module loading, and the top-ranking genes were used for downstream functional annotation. Gene ontology (GO) enrichment analysis was performed using the top module genes relative to the background set of genes included in the cNMF analysis (i.e., expressed genes passing quality-control and feature-selection thresholds). Enriched biological processes were identified using overrepresentation analysis with multiple hypothesis correction, and GO terms were ranked by adjusted p-value.

### Imputation of AT8 levels in a unimodal dataset

Seurat reference mapping was used to transfer AT8 signal from Soma-seq to the Otero-Garcia et al dataset. Both datasets were first subset to L2/3 IT excitatory neurons and re-processed with NormalizeData, FindVariableFeatures(nfeatures = 3000), ScaleData, and RunPCA(npcs = 50) using default parameters. Soma-seq cells were used as the reference, with RNA profiles used to identify anchors and normalized AT8 ADT levels used as the transferred feature. Transfer anchors were identified with FindTransferAnchors(dims=50), and normalized AT8 ADT levels were transferred to the Otero-Garcia query with TransferData(dims = 1:150), both steps run with default parameters. For each sample, mean predicted log-normalized AT8 levels were computed and compared across AT8 FACS groups.

### Protein-protein correlations

To quantify pairwise relationships between intracellular protein markers, we computed pairwise Spearman correlations on the median-normalized ADT counts for 15 epitopes (AT8, HT7, D1M9X, AT180, MAP2, NCAM1, CD142, GFAP, Lamp2, S6, GAD65, p62, ACTB, ms-IgG1, and rb-IgG) across all cells. Epitopes were ordered by hierarchical clustering for visualization.

### Protein-RNA correlations

To assess concordance between protein and transcript abundance at single-cell resolution, each ADT was paired with its cognate RNA (e.g., AT8:MAPT, GFAP:GFAP, p62:SQSTM1; 17 pairs total). Correlations were computed separately within each major cell type (astrocytes, excitatory neurons, inhibitory neurons, and microglia/PVMs) to avoid confounding by cell-type-specific expression differences. Within each cell type, we applied k-nearest-neighbor (KNN) smoothing to reduce single-cell measurement noise. Principal component analysis (PCA) was performed on the RNA expression matrix, and for each cell, its 25 nearest neighbors were identified in the first 20 principal components. Both ADT and RNA values were then averaged across each cell’s local neighborhood to obtain smoothed per-cell estimates. Spearman rank correlations were computed on the smoothed values after excluding cells with zero expression in either modality (minimum 5 non-zero cells). Protein-RNA pairs not biologically relevant to a given cell type (e.g., tau epitopes in astrocytes, glial markers in neurons) were masked from the heatmap.

### iPSC-neuron screen analysis

Aggregation density was calculated as Agg_Count / Dendra_Area for each well. Values were normalized to the mean of AAV2 + seed control wells on the same plate and timepoint, yielding a fold change relative to control. For each perturbation, normalized aggregation density values were pooled across both timepoints (Week 6 and Week 8) and tested against 1.0 (no change vs. control) using a one-sample two-tailed Student’s t-test. P-values were corrected for multiple comparisons using the Benjamini–Hochberg false discovery rate (FDR) method. Z-scores were computed as (mean fold change – 1) / σ, where σ is the standard deviation of the fold change deviations across all perturbations. A gene was considered a significant hit if it met both criteria: FDR-adjusted p < 0.05 and |z-score| > 1.

## Contributions

C.F., C.G., B.A.F., M.B.C conceived the study. C.F. performed all single cell experiments. O.F. and J.H. selected and performed IHC on tissues. C.F., C.G. analyzed all data. C.F., M.A.L., M.B.C. performed iPSC-neuron experiments. M.A.L., M.A.T., C.F., M.B.C analyzed imaging data. C.F., C.G., M.B.C. wrote the manuscript and produced figures. B.A.F., C.C.H., O.R.R., M.B.C. supervised the work. All authors edited the manuscript.

## Extended Data

**Extended Figure 1.**
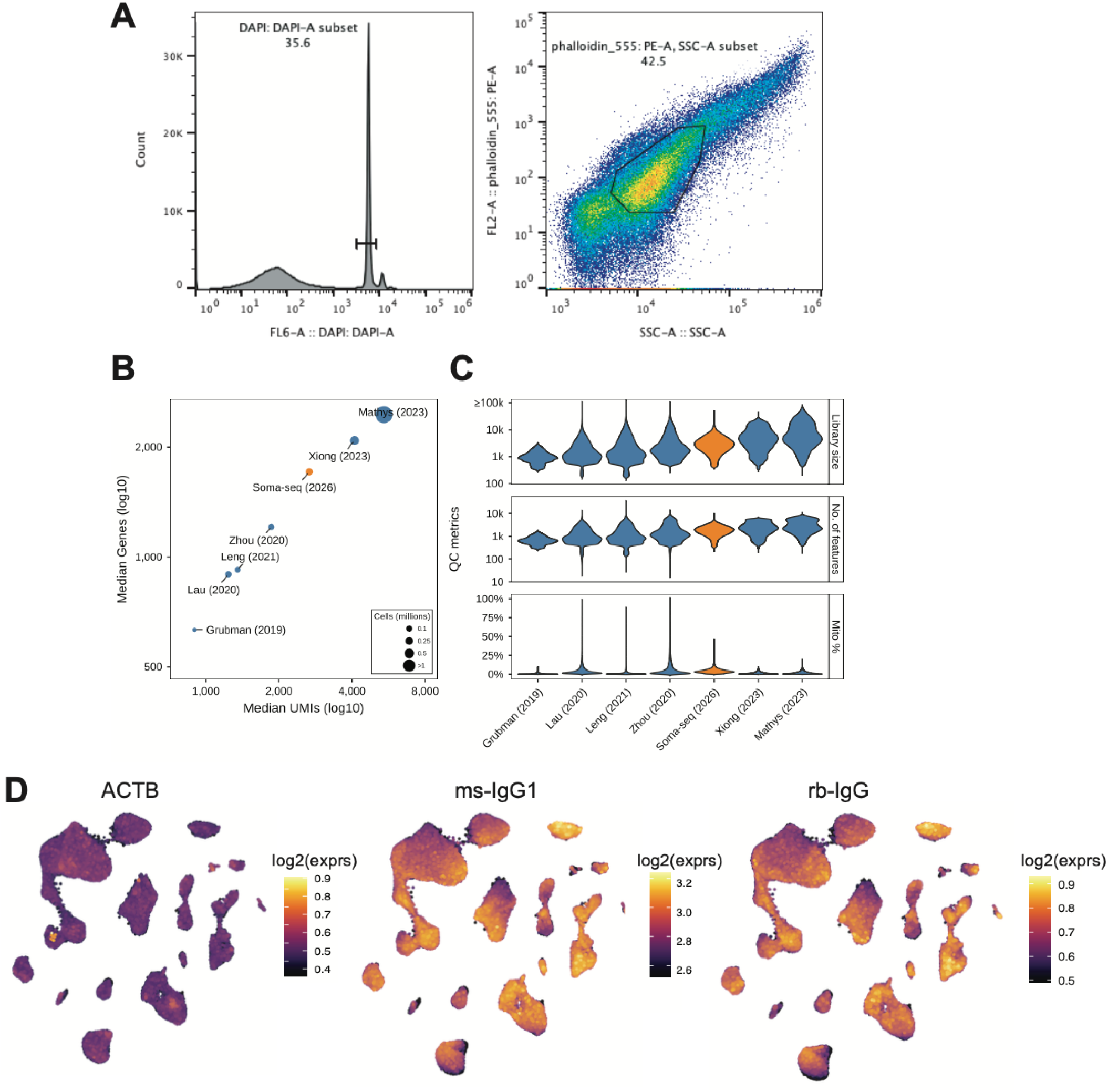
Quality control metrics of Soma-seq. **a**, Example gating strategy for somas: (1) DAPI positive population followed by (2) an intermediate side scatter (area) and phalloidin positive population. **b**, Comparison of median genes and median UMIs detected across published human Alzheimer’s disease brain single-nucleus RNA-seq studies (blue) and Soma-seq (orange). **c**, Comparison of library size, number of detected features, and percentage of mitochondrial transcripts (Mito %) across published human Alzheimer’s disease brain single-nucleus RNA-seq studies (blue) and Soma-seq (orange). **d**, Featureplots of non-cell-type-specific antibody controls, including ACTB, ms-IgG1, and rb-IgG.

**Extended Figure 2.**
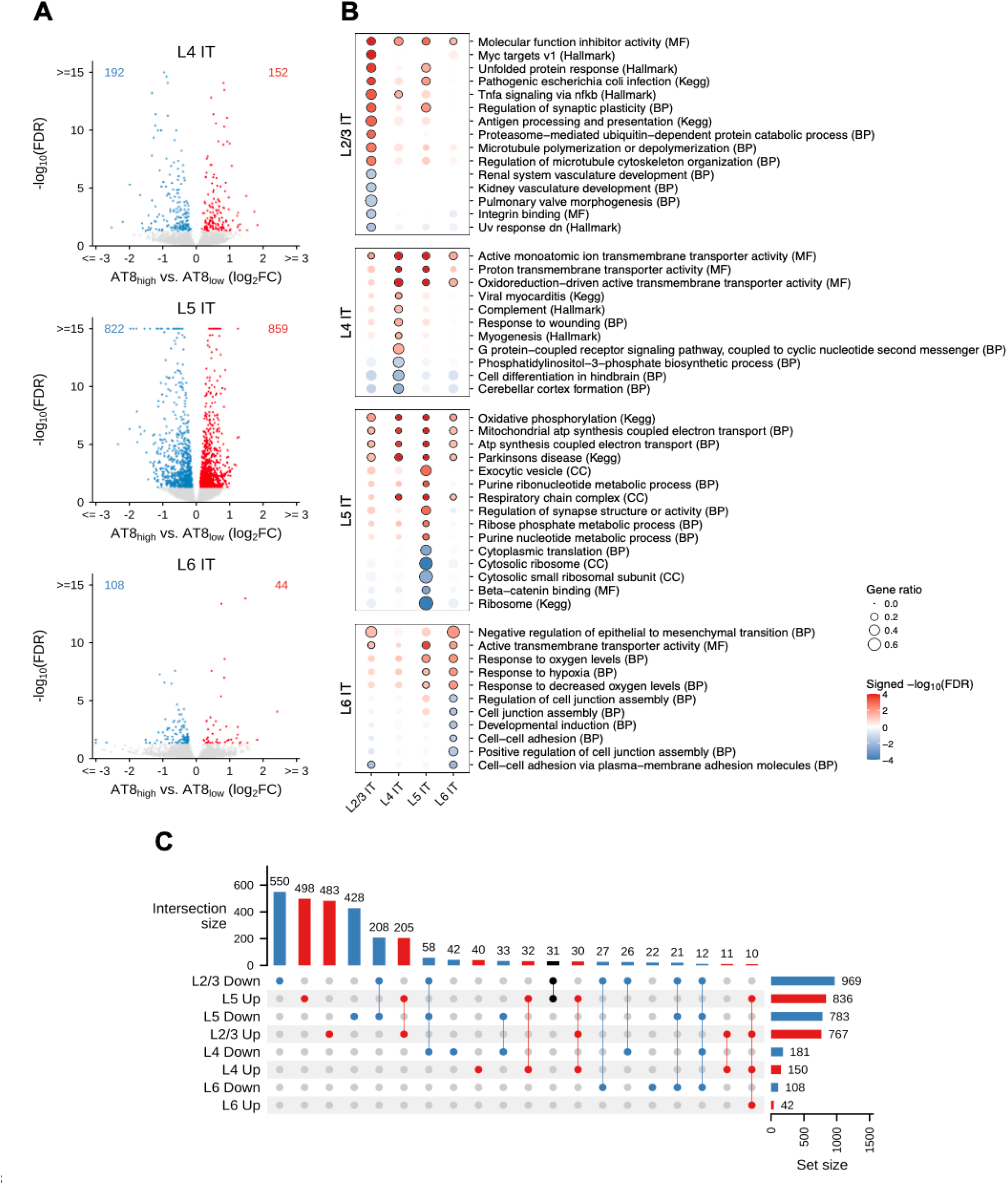
AT8-associated differential gene expression across neuronal layers. **a**, Volcano plots of differentially expressed genes comparing AT8 high and AT8 low neurons across neuronal layers. Red and blue dots indicate genes significantly up- and downregulated, respectively, in AT8 high neurons. FDR ≤ 0.05. **b**, Dot plot of enriched and de-enriched pathways identified from multiple gene set collections, including Biological Process (BP), Molecular Function (MF), Cellular Component (CC), Hallmark, and KEGG, across neuronal layers. Color indicates signed –log10(FDR), and dot size indicates gene ratio. Black outlines denote significance (FDR ≤ 0.05). **c**, UpSet plot depicting the overlap of AT8-associated differentially expressed genes across neuronal layers. Genes upregulated across layers are shown in red, genes downregulated across layers in blue, and genes with opposing directions of regulation across layers in black.

**Extended Figure 3.**
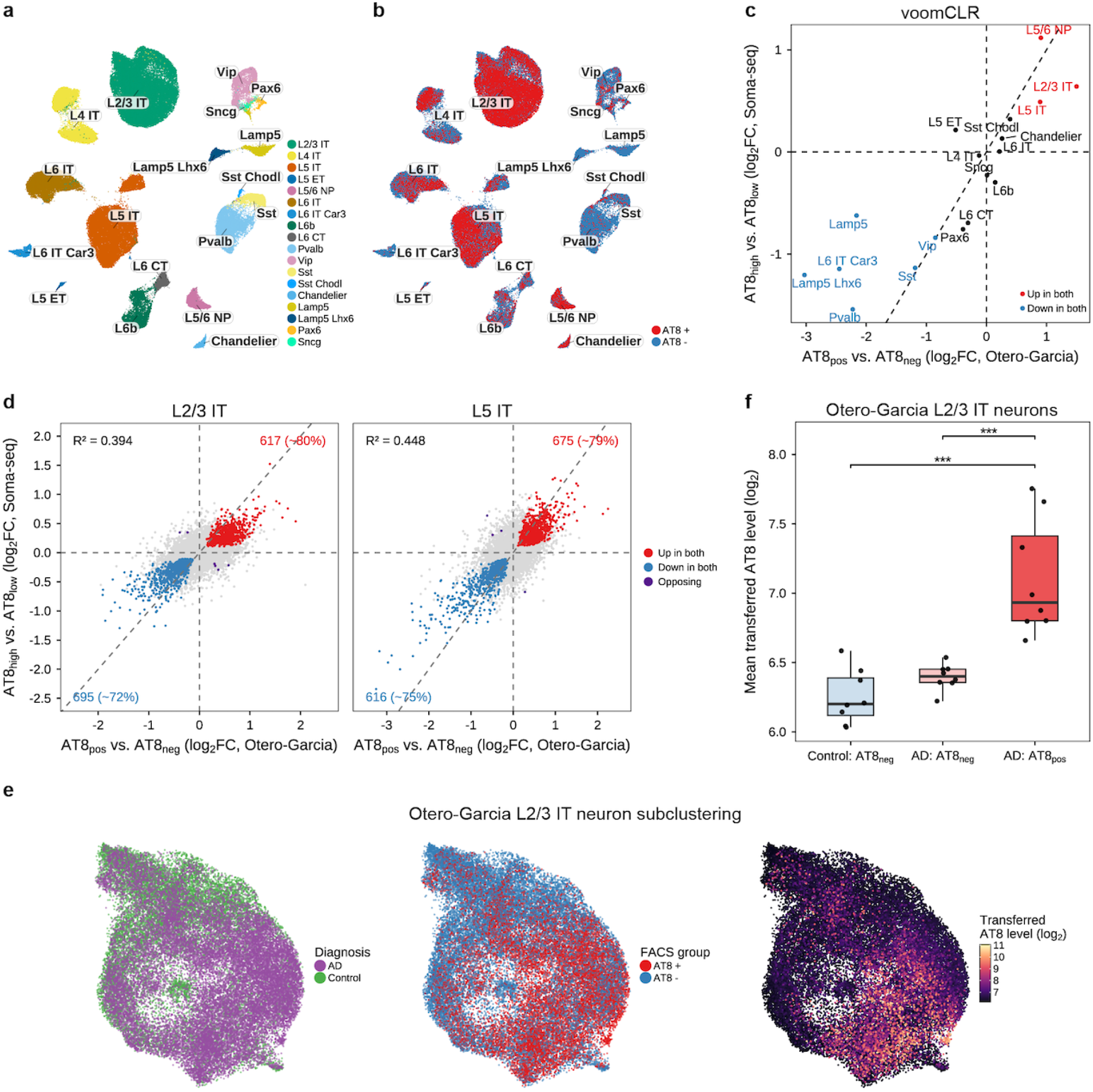
Comparison of AT8-associated changes with Otero et al^**18**^. **a-b**, UMAP representation of the re-analyzed Otero et al. dataset colored by neuronal subtype (**a**) or AT8 FACS gate (**b**). **c**, Comparison of voomCLR-estimated neuronal subtype enrichment and depletion in AT8-high versus AT8-low populations in Soma-seq and AT8-positive versus AT8-negative populations in Otero et al. Red and blue dots indicate neuronal subtypes that are significantly more or less abundant, respectively, in the AT8-high/positive population in both studies. **d**, Scatter plots comparing gene expression changes in AT8-high versus AT8-low neurons in Soma-seq with gene expression changes in AT8-positive versus AT8-negative neurons in Otero et al., shown for L2/3 IT neurons (left) and L5 IT neurons (right). Red and blue dots indicate genes significantly up- and downregulated, respectively, in both datasets. Purple dots indicate genes with opposing directions of regulation. FDR ≤ 0.05. **e**, UMAP representation of sub-clustered L2/3 IT neurons from the Otero-Garcia et al. dataset colored by diagnosis, AT8 FACS gate, or RNA-anchor transferred AT8 level. **f**, Box plots showing mean transferred AT8 levels in L2/3 IT neurons from the re-analyzed Otero et al. dataset, illustrating recovery of known differences in AT8 levels in an independent cohort (see Methods). From left to right, groups correspond to AT8-negative neurons from control subjects (blue), AT8-negative neurons from AD subjects (light red), and AT8-positive neurons from AD subjects (red). Soma-seq AT8 levels in L2/3 IT neurons were used to impute AT8 levels in the Otero et al. dataset. Each dot represents an individual sample from Otero et al. AT8-positive and AT8-negative designations are based on the original FACS sorting strategy in Otero et al.

**Extended Figure 4.**
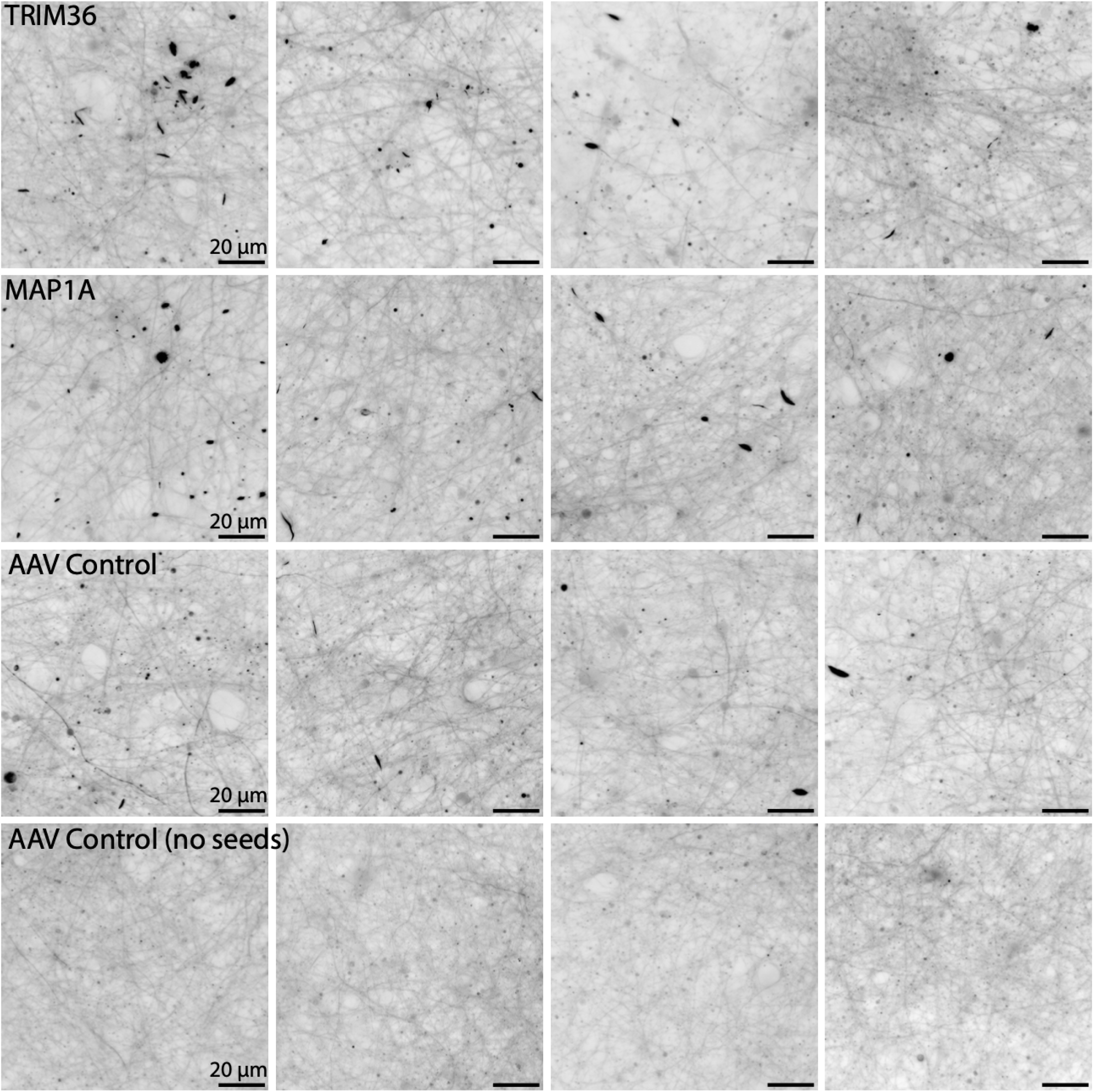
Representative images of Tau aggregates in iPSC-neuron CRISPR screen. Representative images from multiple wells for each gene knockout. Signal represents thresholded Dendra Tau intensity, with punctuated structures indicating Tau aggregates.

## References

1. Morabito, S. et al. Single-nucleus chromatin accessibility and transcriptomic characterization of Alzheimer’s disease. Nat. Genet. 53, 1143–1155 (2021).

2. Leng, K. et al. Molecular characterization of selectively vulnerable neurons in Alzheimer’s disease. Nat. Neurosci. 24, 276–287 (2021).

3. Gabitto, M. I. et al. Integrated multimodal cell atlas of Alzheimer’s disease. Nat. Neurosci. 27, 2366–2383 (2024).

4. Grubman, A. et al. A single-cell atlas of entorhinal cortex from individuals with Alzheimer’s disease reveals cell-type-specific gene expression regulation. Nat. Neurosci. 22, 2087–2097 (2019).

5. Kovacs, G. G. Molecular Pathological Classification of Neurodegenerative Diseases: Turning towards Precision Medicine. International Journal of Molecular Sciences 17, 189 (2016).

6. Wu, T. et al. Single-cell proteomic landscape of the developing human brain. Nature Biotechnology (2026) doi:10.1038/s41587-025-02980-7.

7. Jiang, L. et al. A Quantitative Proteome Map of the Human Body. Cell 183, 269–283.e19 (2020).

8. Ding, Y. et al. Comprehensive human proteome profiles across a 50-year lifespan reveal aging trajectories and signatures. Cell 188, 5763–5784.e26 (2025).

9. Tasaki, S. et al. Inferring protein expression changes from mRNA in Alzheimer’s dementia using deep neural networks. Nature Communications 13, 655 (2022).

10. Braak, H. & Braak, E. Neuropathological stageing of Alzheimer-related changes. Acta Neuropathologica 82, 239–259 (1991).

11. Fujiwara, H. et al. α-Synuclein is phosphorylated in synucleinopathy lesions. Nature Cell Biology 4, 160–164 (2002).

12. Neumann, M. et al. Ubiquitinated TDP-43 in Frontotemporal Lobar Degeneration and Amyotrophic Lateral Sclerosis. Science 314, 130–133 (2006).

13. Stoeckius, M. et al. Simultaneous epitope and transcriptome measurement in single cells. Nature Methods 14, 865–868 (2017).

14. Mimitou, E. P. et al. Scalable, multimodal profiling of chromatin accessibility, gene expression and protein levels in single cells. Nature Biotechnology 39, 1246–1258 (2021).

15. Chung, H. et al. Joint single-cell measurements of nuclear proteins and RNA in vivo. Nature Methods 18, 1204–1212 (2021).

16. Chen, A. F. et al. NEAT-seq: simultaneous profiling of intra-nuclear proteins, chromatin accessibility and gene expression in single cells. Nature Methods 19, 547–553 (2022).

17. Blair, J. D. et al. Phospho-seq: integrated, multi-modal profiling of intracellular protein dynamics in single cells. Nature Communications 16, 1346 (2025).

18. Otero-Garcia, M. et al. Molecular signatures underlying neurofibrillary tangle susceptibility in Alzheimer’s disease. Neuron 110, 2929–2948.e8 (2022).

19. Grubman, A. et al. A single-cell atlas of entorhinal cortex from individuals with Alzheimer’s disease reveals cell-type-specific gene expression regulation. Nat. Neurosci. 22, 2087–2097 (2019).

20. Zhou, Y. et al. Human and mouse single-nucleus transcriptomics reveal TREM2-dependent and TREM2-independent cellular responses in Alzheimer’s disease. Nature Medicine 26, 131–142 (2020).

21. Lau, S.-F., Cao, H., Fu, A. K. Y. & Ip, N. Y. Single-nucleus transcriptome analysis reveals dysregulation of angiogenic endothelial cells and neuroprotective glia in Alzheimer’s disease. Proceedings of the National Academy of Sciences 117, 25800–25809 (2020).

22. Leng, K. et al. Molecular characterization of selectively vulnerable neurons in Alzheimer’s disease. Nat. Neurosci. 24, 276–287 (2021).

23. Morabito, S. et al. Single-nucleus chromatin accessibility and transcriptomic characterization of Alzheimer’s disease. Nat. Genet. 53, 1143–1155 (2021).

24. Xiong, X. et al. Epigenomic dissection of Alzheimer’s disease pinpoints causal variants and reveals epigenome erosion. Cell 186, 4422–4437.e21 (2023).

25. Chen, A. F. et al. NEAT-seq: simultaneous profiling of intra-nuclear proteins, chromatin accessibility and gene expression in single cells. Nature Methods 19, 547–553 (2022).

26. Mimitou, E. P. et al. Scalable, multimodal profiling of chromatin accessibility, gene expression and protein levels in single cells. Nature Biotechnology 39, 1246–1258 (2021).

27. Goedert, M., Jakes, R. & Vanmechelen, E. Monoclonal antibody AT8 recognises tau protein phosphorylated at both serine 202 and threonine 205. Neuroscience Letters 189, 167–170 (1995).

28. Tsaka, G. et al. Conformation-specific monoclonal antibodies reveal early Tau structural intermediates in Alzheimer’s disease. J. Biol. Chem. 302, 111221 (2026).

29. Braak, H. & Tredici, K. D. The preclinical phase of the pathological process underlying sporadic Alzheimer’s disease. Brain 138, 2814–2833 (2015).

30. Nelson, P. T. et al. Correlation of Alzheimer Disease Neuropathologic Changes With Cognitive Status: A Review of the Literature. Journal of Neuropathology & Experimental Neurology 71, 362–381 (2012).

31. Baran, Y. et al. MetaCell: analysis of single-cell RNA-seq data using K-nn graph partitions. Genome Biology 20, 206 (2019).

32. Buerger, K. et al. CSF phosphorylated tau protein correlates with neocortical neurofibrillary pathology in Alzheimer’s disease. Brain 129, 3035–3041 (2006).

33. Pankiv, S. et al. p62/SQSTM1 Binds Directly to Atg8/LC3 to Facilitate Degradation of Ubiquitinated Protein Aggregates by Autophagy*. Journal of Biological Chemistry 282, 24131–24145 (2007).

34. Kotliar, D. et al. Identifying gene expression programs of cell-type identity and cellular activity with single-cell RNA-Seq. eLife 8, e43803 (2019).

35. Zhang, Y. et al. Rapid Single-Step Induction of Functional Neurons from Human Pluripotent Stem Cells. Neuron 78, 785–798 (2013).

36. Croft, C. L. et al. Photodynamic studies reveal rapid formation and appreciable turnover of tau inclusions. Acta Neuropathologica 141, 359–381 (2021).

37. Manos, J. D. et al. Uncovering specificity of endogenous TAU aggregation in a human iPSC-neuron TAU seeding model. iScience 25, 103658 (2022).

38. Mascaro, M., Lages, I. & Meroni, G. Microtubular TRIM36 E3 Ubiquitin Ligase in Embryonic Development and Spermatogenesis. Cells 11, 246 (2022).

39. Halpain, S. & Dehmelt, L. The MAP1 family of microtubule-associated proteins. Genome Biology 7, 224 (2006).

40. Thornburg-Suresh, E. J. C. & Summers, D. W. Microtubules, Membranes, and Movement: New Roles for Stathmin-2 in Axon Integrity. Journal of Neuroscience Research 102, e25382 (2024).

41. Niikura, T., Kita, Y. & Abe, Y. SUMO3 Modification Accelerates the Aggregation of ALS-Linked SOD1 Mutants. PLoS ONE 9, e101080 (2014).

42. Fleming, S. J. et al. Unsupervised removal of systematic background noise from droplet-based single-cell experiments using CellBender. Nat Methods 20, 1323–1335 (2023).

43. Muskovic, W. & Powell, J. E. DropletQC: improved identification of empty droplets and damaged cells in single-cell RNA-seq data. Genome Biol 22, 329 (2021).

44. Heaton, H. et al. Souporcell: robust clustering of single-cell RNA-seq data by genotype without reference genotypes. Nat Methods 17, 615–620 (2020).

45. Lun, A. T. L. et al. EmptyDrops: distinguishing cells from empty droplets in droplet-based single-cell RNA sequencing data. Genome Biol 20, 63 (2019).

46. Germain, P.-L., Lun, A., Meixide, C. G., Macnair, W. & Robinson, M. D. Doublet identification in single-cell sequencing data using scDblFinder. F1000Res 10, 979 (2021).

47. Gabitto, M. I. et al. Integrated multimodal cell atlas of Alzheimer’s disease. Nat. Neurosci. 27, 2366–2383 (2024).

48. Hao, Y. et al. Dictionary learning for integrative, multimodal and scalable single-cell analysis. Nat Biotechnol 42, 293–304 (2024).

49. Assefa, A. T., Verbist, B. & Berge, K. V. den. Assessing differential cell composition in single-cell studies using voomCLR. Bioinformatics 42, (2026).

